# Transcriptomic divergence predicts morphological and ecological variation underlying an adaptive radiation

**DOI:** 10.1101/2020.12.15.422975

**Authors:** Moisés A. Bernal, Daniel L. Yule, Wendylee Stott, Lori Evrard, Thomas E. Dowling, Trevor J. Krabbenhoft

## Abstract

Groups of sympatric taxa with low inter-specific genetic differentiation, but considerable ecological differences, offer great opportunities to study the dynamics of divergence and speciation. This is the case of ciscoes (*Coregonus* spp.) in the Laurentian Great Lakes, which are characterized by a complex evolutionary history and are commonly described as having undergone an adaptive radiation. In this study, morphometrics, stable isotopes and transcriptome sequencing were used to study the relationships within the *Coregonus artedi* complex in western Lake Superior. We observed general concordance for morphological, ecological and genomic variation, but the latter was more taxonomically informative as it showed less overlap among species in multivariate space. Low levels of genetic differentiation were observed between individuals morphologically identified as *C. hoyi* and *C. zenithicus*, and we hypothesize this could be associated with recent hybridization between the two species. Transcriptome-based single nucleotide polymorphisms exhibited significant divergence for genes associated with vision, development, metabolism and immunity, among species that occupy different habitats. This study highlights the importance of using an integrative approach when studying groups of taxa with a complex evolutionary history, as individual-level analyses of multiple independent datasets can provide a clearer picture of the patterns and processes associated with the origins of biodiversity.

## INTRODUCTION

The speciation continuum offers exciting opportunities to understand the mechanisms driving the accumulation of genetic and ecological divergence among closely related groups. This is especially relevant when ecological opportunities can promote and/or maintain differentiation among such groups, even in the presence of gene flow (Arnold, 2006; Fitzpatrick, Gerberich, Kronenberger, Angeloni, & Funk, 2015). Hybridization during early stages of differentiation can complicate the identification of divergent groups, as selectively neutral alleles may be freely exchanged between forms (Hohenloe et al., 2013; Feulner & Seehausen, 2019). Conversely, genes associated with traits of reproductive or ecological advantage (Hench, Vargas, Höppner, McMillan, & Puebla, 2019; Meier, Marques, Wagner, Excoffier, & Seehausen, 2018; Richards & Martin, 2017) are expected to be far more resistant to gene flow among groups. This scenario of divergence can be further complicated by sporadic or cyclical environmental disturbances, which can disrupt the ecological barriers maintaining isolation between closely related forms (Feulner & Seehausen, 2019; Ohlberger, Mehner, Staaks, & Hölker, 2008), or affect the geographic barriers that hinder gene flow (Turgeon & Bernatchez, 2003; Wilson & Bernatchez, 1998). These patterns are commonly observed in lacustrine environments, such as the Laurentian Great Lakes (hereafter the Great Lakes), where multiple closely related groups/forms occur in sympatry (e.g., *Coregonus* spp., *Salvelinus* spp.).

Aquatic organisms in the Great Lakes are characterized by a complex evolutionary history as a result of Pleistocene glaciation events. Glaciers covered most areas currently associated with the Great Lakes during the Pleistocene, leading to the isolation of lakes that remained ice-free (Bailey & Smith, 1981). The retreat of the ice sheets, which started approximately 14,000 years before present, led to the formation of lakes and rivers that promoted connectivity between the isolated refugia (Bailey & Smith, 1981). These cycles of vicariance followed by subsequent hypothesized admixture resulted in fish groups has led to complex phenotypic and genomic patterns and complicated taxonomic relationships (Bernatchez & Dodson, 1991; Turgeon, Estoup, & Bernatchez, 1999; Wilson & Bernatchez, 1998). One example is the ciscoes in the genus *Coregonus* (subgenus *Leucichthys*), a group that contains several species characterized by distinct morphological and ecological traits. An early study of this group recognized eight distinct Great Lakes species in the *C. artedi* complex: *C. alpenae*, *C. artedi*, *C. johannae*, *C. kiyi*, *C. hoyi*, *C. nigripinnis*, *C. reighardi* and *C. zenithicus*, as well as several sub-species (Koelz, 1929). Later work has resulted in considerable debate over the systematics of the group, due to phenotypic and genetic similarity among some of the species (Todd, Smith, & Cable, 1981; Turgeon & Bernatchez, 2003; Turgeon, Estoup, & Bernatchez, 1999).

During the 20^th^ century, human mediated effects such as overfishing, pollution, habitat degradation and invasive species directly affected the Great Lakes (Mills, Leach, Carlton, & Secor, 1994; Regier, Whillans, Christie, & Bocking, 1999). These changes led to population declines and extinctions of many coregonines (Smith, 1968). For example, it is thought that overfishing led to the extinction of six deep-water cisco forms in Lake Michigan (Eshenroder et al., 2016). Moreover, there has been a substantial decline in the abundance of deep-water forms in Lake Superior since the 1920s, especially *C. zenithicus* (Bronte et al., 2010; Hoff & Todd, 2004; Pratt, 2012). In many instances, population declines and extinctions have been followed by expansion of more abundant species into habitats previously occupied by other groups. Such is the case for *C. hoyi* which now occurs in deep waters that were once occupied by other species in lakes Michigan and Huron (Bunnell et al., 2012). Studies in marine and freshwater fishes have shown that habitat degradation and other types of environmental change, along with fluctuations in population density can lead to blurring of ecological and behavioral barriers that keep closely related groups separated when they lack Seehausen, 2006; Taylor et al., 2006). Over time, the erosion of pre-mating mechanisms can lead to loss of taxonomic diversity via “speciation reversal” (Zhang, Thibert-Plante, Ripa, Svanbäck, & Brännström, 2019). It has also been suggested that reduction in population densities for one taxon can lead to increased rates of hybridization with other more abundant and closely related groups (i.e., Hubbs principle; Hubbs, 1955), as it becomes challenging for the rare species to find mates of their own kind. Hybridization can quickly exacerbate the loss of genetic diversity caused by the initial population declines (Arnold, Bulger, Burke, Hempel, & Williams, 1999). Previous studies have suggested that hybridization between closely related groups could have accelerated the extinction of cisco forms, representing a loss of biodiversity (Todd & Stedman, 1989). Given the sympatric nature of Lake Superior ciscoes and their low levels of genetic differentiation (Ackiss, Larson, & Stott, 2020; Turgeon & Bernatchez 2003; Turgeon, Estoup, & Bernatchez, 1999), it is important to understand the phenotypic and genetic differences that characterize the extant members of the *C. artedi* complex, to improve the understanding of their evolutionary history and to inform conservation efforts in Lake Superior.

Today, up to six species are thought to be present in Lake Superior (Eshenroder et al., 2016), and three are routinely caught with standard trawl or gillnet sampling: *C. artedi, C. hoyi* and *C. kiyi* (US Geological Survey, 2018). The most abundant is *C. kiyi* (Yule et al., 2013) and it tends to occupy bathymetric depths greater than 100m (Rosinski, Vinson, & Yule, 2020) and predominantly feeds on *Mysis diluviana* (Gamble, Hrabik, Stockwell, & Yule, 2011a; Ahrenstorff et al., 2011). The second most abundant form is *C. artedi* (Yule et al., 2013) which is found near the lake surface, especially during spring and summer (Stockwell, Yule, Gorman, Isaac, & Moore, 2006; Yule et al., 2013; Rosinski, Vinson, & Yule, 2020). The primary diet of adult *C. artedi* consists of *Mysis*, cladocerans and calanoid copepods (Gamble, Hrabik, Stockwell, & Yule, 2011a, b; Ahrenstorff, Hrabik, Stockwell, Yule & Sass, 2011). *Coregonus hoyi* is the third most abundant form and can be associated with the lakebed during the day and found in waters of the hypolimnion at night (Yule et al., 2013). They primarily consume calanoid copepods and cladocerans as juveniles, while adults utilize a diverse diet of zooplankton, *Diporeia* spp. and *Mysis* (Gamble, Hrabik, Stockwell, & Yule, 2011b; Sierszen et al., 2014). *Coregonus zenithicus* is also extant but is thought to be extremely rare (Bronte et al., 2010; Hoff & Todd, 2004; but see Pratt, 2012), while the status of *C. nigripinnis* and *C. reighardi* is uncertain (Eshenroder et al, 2016; Todd & Smith, 1980). Previous studies in Lake Superior using stable isotopes over three time-frames (1897-1929, 1934-1966, and 1972-1998) concluded that *C. zenithicus* and *C. nigripinnis* had a high degree of overlap in both trophic position and their use of basal carbon resources, while *C. reighardi* was distinct from these other two species (Schmidt et al., 2009). Overall, it is estimated that *C. zenithicus* consumes primarily *Mysis*, *Diporeia* spp., calanoid copepods and cladocerans (reviewed by Eshenroder et al., 2016). However, due to their rarity, it has not been possible to include them in more recent studies.

Due to their considerable ecological differences, the *Coregonus artedi* species complex provides a unique opportunity to explore mechanisms that lead to the differentiation and maintenance of closely related sympatric forms. There have been many previous genetic studies of the *Coregonus* species complex in the Great Lakes, and most have relied on putatively neutral markers (i.e., mitochondrial DNA and microsatellites). Some studies have identified significant differences associated with geographic regions but have failed to identify differences among species (Turgeon & Bernatchez, 2003; Turgeon, Estoup, & Bernatchez, 1999). A more recent study used single nucleotide polymorphisms (SNPs) generated by RADseq to identify genetic differentiation among forms in Lake Superior, but did not include morphological or ecological data (Ackiss, Larson, & Stott, 2020). Since morphological and ecological divergence is partially heritable in this group (Todd & Smith, 1980), we anticipate that there is underlying functional genetic divergence in some parts of the genome, which could be identified by comparing the transcriptomes of species in the *Coregonus artedi* complex. An initial study examined functional differences in opsin genes across the complex, and found evidence of adaptive molecular evolution in *rhodopsin*, consistent with differences in depth preferences among forms (Eaton et al. 2020), suggesting that functional genetic differences might also be present in other gene families.

In this study, we aim to understand if there is concordance between morphology, feeding habits, and genomic divergence among four species of *Coregonus* from Lake Superior: *C. artedi*, *C. kiyi*, *C. hoyi*, and *C. zenithicus*. Despite ongoing debates on whether *C. nigripinnis* and *C. reighardi* should be synonymized with *C. zenithicus* (Eshenroder et al., 2016; Todd & Smith, 1980), these groups were not included in the present study as they are very rare in Lake Superior. Specific questions for our study include: 1) What are the main morphological traits that characterize different species? 2) What are the main ecological traits that differentiate the deep-water ciscoes *C. hoyi* and *C. zenithicus*? 3) Which highly divergent genes are associated with ecological and morphological differences? By understanding the phenotypic, ecological, and genomic differences among these species, this study presents an integrative view of the divergence of fishes of the Laurentian Great Lakes.

## METHODS

### Sample collections

Individuals of *C. artedi, C. hoyi, C. kiyi* and *C. zenithicus* were collected between May and November of 2015 from multiple sites throughout western Lake Superior (Figure 1, Table S1). Identification of individuals followed Koelz (1929) and Eshenroder et al. (2016). Extant forms of *C. hoyi* and *C. zenithicus* were identified based on length of the lower jaw (lower jaw is shorter for *C. zenithicus*; Eshenroder et al., 2016) and a cursory examination of the number of gill-rakers (*C. zenithicus* has 35-40 and *C. hoyi* 41-45; Eshenroder et al., 2016). Captured fish were identified to species, euthanized by pithing, and dissected to obtain tissues for RNA isolation. In total, eight individuals per species were included in all analyses, except for *C. zenithicus* (n=7; see Results). All sampling and handling of fish was done following the guidelines for the care and use of fishes by the American Fisheries Society (AFS 2014).

**Figure 1.**
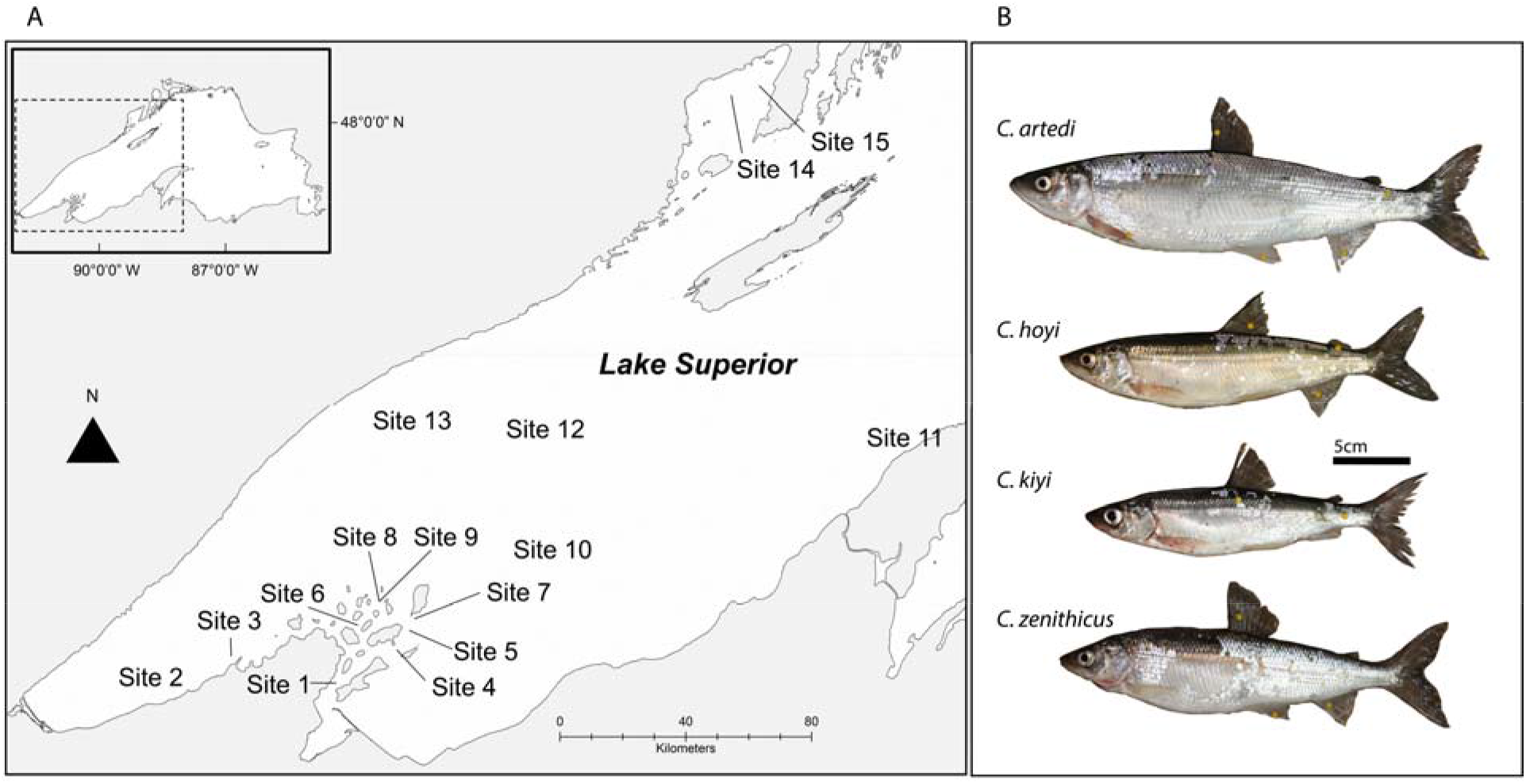
(A) Collection sites for organisms analyzed in the study and (B) sympatric forms of Lake Superior coregonines: *Coregonus artedi*, *C. hoyi*, *C. kiyi* and *C. zenithicus*. Detailed information about sites are provided in Supplementary Table S1. Photographs by D.L Yule.

### Morphometrics

After dissection for tissues that would be used in RNAseq analyses, fish were placed on ice until they could be photographed. Fishes were placed on a mesh on top of a wooden frame with a ruler for scale (mm). Attempts were made to position specimens in a natural position, with the mouth slightly opened (Eshenroder et al., 2016). Dorsal, adipose, caudal and anal fins were extended with pins to allow for their measurement. Pelvic and pectoral fins were positioned so the horizontal and distal ends were clearly visible. A card with the specimen number and species field identification was included in each photograph. A digital image of the whole body was taken, and each image was scrutinized to ensure that all landmarks needed for body measurements were visible. Individuals were then frozen at −20°C to preserve tissues for the stable isotope analysis.

Digital images were used to obtain 11 body measurements described in Eshenroder et al. (2016): body depth (BDD), dorsal fin height (DOH), head length (HLL), maxillary length (MXL), orbital length (OOL), pelvic-anal distance (PAD), pectoral fin length (PCL), preorbital length (POL), pelvic fin length (PVL), pectoral-pelvic distance (PPD), and standard length (STL; Table S2). The lengths of the PCL and PVL were divided by PPD and PAD, respectively, providing fin-length-to-body-distance ratios. These ratios have been shown to be useful for distinguishing *Coregonus* species (Koelz, 1929; Eshenroder et al., 2016). The premaxillary angle (PMA) was also measured with a protractor but was not included in the PCA analysis because a cursory analysis showed PMA was taxonomically uninformative.

The image analysis software SigmaScan Pro V5.0.0 was used to obtain all measurements, calibrating the distances with the ruler included in each image. Body measurements (in mm) were transformed to their natural log. A principal component analysis (PCA) was used to capture the maximum amount of variation in just a few dimensions, using the *prcomp* function in R, with default settings (R Core Team 2013). To remove the size component from the morphometric measures, the covariance matrix was estimated, and the first principal component of these measures was used as a general measure of size (Dos Reis, Pessôa, & Strauss, 1990). Except for the two fin ratios, all transformed measurements were regressed against the first principal component to obtain size-free residuals that we analyzed in a second PCA.

### Stable Isotope Analysis

We used stable isotope ratios (δ^13^C and δ^15^N) to estimate isotopic niches and trophic overlap of the identified forms. Individuals stored at −20°C were thawed and white muscle tissue was removed posterior to the dorsal fin. Skin was removed and the sample rinsed in deionized water, dried at 50-60°C, homogenized, and 0.5-1.0mg was packed into tin capsules. Samples were analyzed at the University of California—Davis Stable Isotope Facility (http://stableisotopefacility.ucdavis.edu/). To accurately interpret the δ^13^C values it is important to normalize for lipid content, especially when C:N_bulk_ > 4 (Hoffman, Sierszen, & Cotter 2015). We applied the arithmetic-mass-balance normalization of Hoffman, Sierszen, & Cotter (2015) to all samples: δ^13^C_lipid free_ = δ^13^C_bulk_ + (Δδ^13^C_lipid_ * (C:N_lipid free_ – C:N_bulk_)) / C:N_bulk_. For this equation, C:N_bulk_ was molar, C:N_lipid free_= 3.8 and the lipid discrimination (Δδ^13^C_lipid_)= −6.4‰. The δ^15^N values were not corrected to allow for detection of δ^15^N enrichment, which occurs with increasing depth in Lake Superior benthos but not zooplankton (Sierszen et al., 2006; 2014).

Overlap of isotopic niches (*sensu* Newsome, Martinez del Rio, Bearhop, & Phillips, 2007) was examined using Stable Isotope Bayesian Ellipses, in R-SIBER v2.1.3 (Jackson, Inger, Parnell, & Bearhops, 2011). Corrected standard ellipse areas (SEAc) were developed so that a subsequently sampled data point would have a 95% probability of being encompassed by a given SEAc. This was accomplished by multiplying the semi-major and -minor axes by 2.45 (Jackson, Inger, Parnell, & Bearhop, 2011). The percentage of overlap between form ellipses were calculated by using the *maxLikOverlap* function in R (R Core Team, 2013).

### Transcriptome sequencing, assembly and SNP analysis

For the transcriptomic analyses, we collected tissues that are expected to play important roles in functional differentiation among *Coregonu*s species: the right half of upper and lower jaws and associated muscle and skin, first gill arch, eye, and brain. Tissue samples were obtained from the same individuals used in the morphological and dietary analyses immediately after pithing. These tissues are actively growing in adult fish, and therefore are expected to show variation based on morphological and ecological differences of the analyzed species. Collected tissues were preserved in RNAlater (Ambion, Inc.) and similar amounts were homogenized and pooled for the RNA extractions. Total RNA was extracted from each homogenate using TRIzol (Life Technologies, Inc.) and RNA Mini Isolation Kits (Ambion, Inc.) following manufacturers’ instructions (including treatment with DNase). RNA quality was assessed with a Bioanalyzer 2100 and quantified using a Qubit 2.0 fluorometer (Life Technologies, Inc.). Library preparation was done at the RTSF Genomics Core facility at Michigan State University, using Illumina TruSeq RNA library kits, with Illumina Ribo-Zero Gold for rRNA depletion. Libraries were normalized and pooled for multiplex sequencing, and the pool was loaded on both lanes of an Illumina HiSeq 2500 Rapid Run flow cell (paired-end, 250bp). Removal of Illumina adaptors and quality filtering (Q > 30) was done with TrimGalore (Krueger, 2015). Resulting sequences were used to assemble a reference transcriptome for each species using Trinity (Grabherr et al., 2011), and only contigs longer than 300bp were maintained. Contigs were summarized using Transdecoder (Haas et al., 2013) which only retains samples with predicted open reading frames. CDHIT (Li & Godzik, 2006) was used to cluster contigs by similarity (95%), matching to the most similar cluster in the dataset. The completeness of *C. artedi* transcriptomes (Trinity *vs.* summarizing with Transdecoder and CDHIT) were assessed with the Benchmarking Universal Single Copy Ortholog database version 3 (BUSCO; Simão, Waterhouse, Ioannidis, Kriventseva, & Zdobnov, 2015), using the Actinopterygii database (*actinopterygii_odb9*). This resulted in 4,049 complete BUSCOs (874 single-copy and 3,175 duplicated) and 200 missing for the Trinity/CDHIT transcriptome, and 3,876 complete BUSCOs (977 single-copy and 2899 duplicated) and 303 missing BUSCOs for the Transdecoder transcriptome (Figure S1). Given that the Transdecoder approach greatly reduced the redundancy of reads while only minimally reducing the completeness of the transcriptome, this version was chosen for downstream analyses. Annotation of the four transcriptomes was done with BLAST search algorithm, using the Uniprot (accessed January 2019) and Trembl (accessed January 2019) databases as reference and an e-value of 1e-10 and below as cutoff. Additionally, transcripts were annotated using Blastn searches against cDNA sequences from six related fish genomes in Ensembl 99 (Ensembl.org): Danio rerio, *Esox lucius, Hucho hucho, Salmo salar, Salmo trutta*. Default search parameters were used for Blastn searches except -evalue 1e-5 and -max_target_seqs 20. Blast hits with <70% similarity and <100 bp alignment were removed and the hit with the highest HSP of the remaining alignments was retained. Annotation to gene ontology (GO) was obtained with the UniprotKB Retrieve/IDmapping (available at: https://www.uniprot.org/uploadlists/), using the UniProt and Trembl gene names as query and exporting a table with the multiple GO terms per gene name.

To compare sequence variation within and among species, we called single nucleotide polymorphisms (SNPs) in the sequenced protein coding loci or adjacent untranslated regions (UTRs) for each individual. Paired-end sequences of all individuals were mapped to the transcriptome of *C. artedi*, using Bowtie2 (Langmead & Salzberg, 2012) with default parameters. The rate of mapping to the *C. artedi* transcriptome was >82% for all individuals. SNP calling was done using the BCFtools platform (Li et al., 2009), with the commands “*mpileup*” (-a 1, -C50, -Q25), and “*call*” (only keeping variant sites, excluding indels). The accessory script *vcfutils.pl* was used to retain sites with coverage higher than 15x, and *VCFtools* (Danecek et al., 2011) was used to keep SNPs separated by 1,000bp, that were present in all individuals. This resulted in a matrix of 22,285 variant SNPs, with no missing data.

Observed (*H*_*o*_) and expected (*H*_*e*_) heterozygosity, number of alleles (*N*_*a*_), inbreeding coefficient (*F*_*is*_), and genetic diversity (θ) were estimated with “*basic.stats*” of the R package *Pegas* (Paradis, 2010) to quantify levels of genetic variation. The package *Genodive* (Meirmans & Tienderen, 2004) was used to estimate the Fixation Index (*F*_*st*_) for pairwise comparisons between the different species. This approach includes the Bonferroni correction of *p*-values for multiple comparisons (5,000 iterations). *VCFtools* was used to estimate pairwise *F*_*st*_ by individual loci between the different groups.

In addition, two separate analyses were performed to assess the degree of concordance between genomic markers and the results of the morphological analyses. First, a PCA was performed with the R package *pcadapt* (Luu, Bazin, & Blum, 2017), using the transcriptome covariance matrix and compared to PCA results from the morphological data. Second, individuals were divided into groups based on their ancestry using a maximum likelihood approach with the program *Ohana* (Cheng, Mailund, & Nielsen, 2016). The analysis was run for 100 iterations (stable likelihoods were observed after 23 iterations), and resulting estimates were used to generate a population assignment plot using Python scripts included in *Ohana*. To determine if these lineages were genetically admixed, we implemented the three-population analysis (*f3 statistics*; Reich, Thangaraj, Patterson, Price, & Singh, 2009) using Treemix (Pickrell et al., 2012). Here, the genetic composition of one coregonine species was compared to two others, and admixture was implied if the z-score was negative (Reich, Thangaraj, Patterson, Price, & Singh, 2009).

The program *Ohana* (Cheng, Racimo, & Nielsen, 2019) was also used to determine if some of the observed SNPs may be under divergent selection among species. Potential candidate loci were considered to be under selection when the maximum likelihood algorithm detects large differences in allele frequencies among species (i.e., Log-likelihood ratios >10; Cheng, Racimo, & Nielsen, 2019). Individual candidate loci that exhibited elevated *F*_*st*_ and/or were identified as evolving under divergent selection with *Ohana* were examined to determine whether any GO categories were significantly overrepresented among highly divergent genes using a Mann-Whitney U test of ranks (GO-MWU) with Benjamini-Hochberg correction (Wright, Aglyamova, Meyer, & Matz, 2015; scripts available: https://github.com/z0on/GO_MWU). This analysis was performed for the highly differentiated genes between populations, as well as the genes that may be under divergent selection for all species, detected with *Ohana*. This test determines if there is a significant enrichment of GO categories among the top values of the distribution (i.e., highest *F*_*st*_). Only categories that are represented by five or more transcripts were taken into consideration for the analyses, and the cutoff for false discovery rate was 10%. For each comparison, the enrichment was done separately for the domains Biological Process (BP), Cellular Component (CC) and Molecular Function (MF).

Because salmonids underwent an ancestral whole genome duplication ~88 Ma (Lien et al., 2016), there is a possibility that some of the identified SNPs are located in paralogous rather than orthologous loci (i.e., ohnologs). This could inflate estimates of divergence between species. In order to confirm the validity of the results, we filtered the transcripts to only include putatively orthologous sequences between the four target groups of our study and the more distantly related European whitefish *C. lavaretus* (NCBI SRA: SRR6321817-SRR6321824; Carruthers et al., 2018) using *Orthofinder* (Emms & Kelly, 2015). The resulting SNP matrix was used to re-estimate the pairwise *F*_*st*_ following the steps described above.

## RESULTS

One individual of *C. zenithicus* always clustered with samples of *C. kiyi* for both genetic and morphological traits (eye diameter in particular; Figure S2). Upon further examination of the photographs, it was determined this individual was a *C. kiyi*, and due to this initial misidentification, this individual was excluded from the results below to avoid any confusion.

### Morphology

Great Lakes coregonines are challenging to identify based on morphology. The PCA showed that variables with greatest influence on PC1 (Figure 2a; Table 1) were paired fin lengths (PCL and PVL), paired-fin length to body length ratios (PCL/PPD & PVL/PAD), and eye orbit size (OOL). *Coregonus kiyi* had the longest paired fins, *C. artedi* had the shortest, and *C. hoyi* and *C. zenithicus* were intermediate (Table 1). PCA scores clearly separated samples of *C. kiyi* and *C. artedi* and were most influenced by PVL/PAD, PCL and BDD. The morphometric analysis could not differentiate samples of *C. zenithicus* and *C. hoyi* included in our study, as they showed considerable overlap for the evaluated traits (Figure 2a).

**Table 1.** Summary of morphological measurements that were most useful for the differentiation of *Coregonus artedi*, *C. hoyi*, *C. kiyi* and *C. zenithicus*. Values in the table represent the average measures or ratios (±SD). Measurement abbreviations are defined in Supplementary Table S2. The Principal Component Analysis loadings for PC1 and PC2 that exceeded ± 0.2 for one (or both) of the dimensions are also provided. Table abbreviations: PCL=Pectoral fin length, PPD=Pectoral-pelvic distance, PVL=Pelvic fin length, PAD=Pelvic-anal distance, OOL=Orbital length (eye), BDD=Body depth.

|  | Total length (mm) | PCL (mm) | PPD (mm) | PCL/PPD | PVL (mm) | PAD (mm) | PVL/PAD | OOL (mm) | BDD (mm) |
| --- | --- | --- | --- | --- | --- | --- | --- | --- | --- |
| *PC1* | - | 0.6 | -0.22 | 0.47 | 0.25 | -0.15 | 0.4 | 0.24 | -0.14 |
| *PC2* | - | 0.4 | -0.21 | 0.07 | -0.19 | 0.24 | -0.73 | 0.14 | -0.32 |
| *C. artedi* | 332 (21) | 41.0 (3.4) | 87.6 (5.2) | 0.47 (0.05) | 37.1 (2.5) | 63.2 (3.7) | 0.59 (0.03) | 12.6 (0.9) | 64.9 (3.7) |
| *C. hoyi* | 243 (25) | 32.4 (4.1) | 64.0 (6.7) | 0.51 (0.07) | 30.5 (3.2) | 42.2 (5.3) | 0.73 (0.07) | 11.3 (0.6) | 45.6 (5.5) |
| *C. zenithicus* | 261 (20) | 31.9 (5.0) | 68.9 (5.6) | 0.46 (0.07) | 32.1 (3.4) | 46.8 (4.5) | 0.69 (0.07) | 11.5 (0.7) | 51.3 (4.5) |
| *C. kiyi* | 225 (16) | 35.9 (3.3) | 56.2 (4.2) | 0.64 (0.03) | 31.0 (3.4) | 38.1 (4.4) | 0.82 (0.04) | 12.6 (0.7) | 40.2 (5.7) |

**Figure 2.**
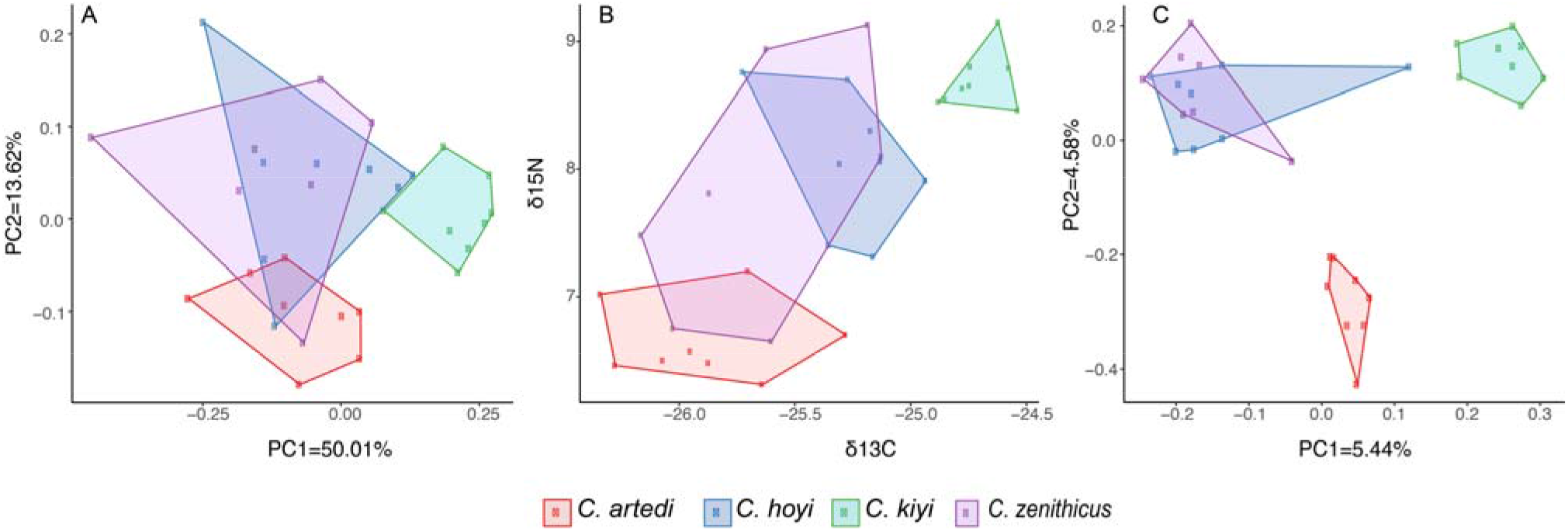
Principal Component Analysis (PCA) for (A) morphological measures; (B) bi-plot for isotope ratios; and (C) PCA of transcriptomic variance (22,285 SNPs) for four species of *Coregonus* from Lake Superior. *Coregonus artedi* (red), *C. hoyi* (blue), *C. kiyi* (green) and *C. zenithicus* (purple).

### Isotopic Niche Breadth and Trophic Overlap

Stable isotope measurements made it possible to differentiate among most of the species (Figure 2b). There was clear separation of convex hulls in isotopic space between the shallow-water *C. artedi* and deep-water *C. kiyi* (Figure 2b, Table 2). Samples of *C. kiyi* generally had the highest mean δ^13^C and δ^15^N values, but a small δ^13^C range (0.3) and δ^15^N range (0.7), indicating this species had the narrowest isotopic niche (Table 2). Meanwhile, samples of *C. artedi* had the lowest mean δ^13^C and δ^15^N values (Table 2). Individuals identified as *C. hoyi* fell between *C. artedi* and *C. kiyi* in the biplot (Figure 2b). These results corroborate previous analyses that suggest *C. hoyi* has an opportunistic feeding strategy and feeds at intermediate depths (Sierszen et al., 2014, Rosinski et al., 2020). Our results showed that samples of *C. zenithicus* overlapped with *C. artedi* and *C. hoyi*.

**Table 2.** The isotopic metrics of δ^13^C_lipid free_ and δ^15^N (‰), for *C. artedi*, *C. hoyi*, *C. kiyi* and *C. zenithicus* collected from western Lake Superior (May-November 2015), and the corrected standard ellipse areas (SEAc).

| | Sample size | Mean $\delta^{15}\text{N}$ (range) | Mean $\delta^{13}\text{C}$ (range) | SEAc |
| --- | --- | --- | --- | --- |
| *C. artedi* | 8 | 6.7 (0.9) | -25.9 (1.1) | 2.33 |
| *C. hoyi* | 8 | 8.1 (1.5) | -25.3 (0.8) | 2.41 |
| *C. zenithicus* | 7 | 7.8 (2.5) | -25.7 (1.0) | 7.19 |
| *C. kiyi* | 8 | 8.7 (0.7) | -24.7 (0.3) | 0.57 |

The SEAc were smallest for *C. kiyi*, intermediate for *C. artedi* and *C. hoyi* and largest for *C. zenithicus* (Table 2; Figure S3). There was no overlap between the ellipses representing *C. artedi* and *C. kiyi*, and there was minimal overlap between these two species and *C. hoyi* (Table 3). This supports a scenario of strong niche partitioning between these three species. In sharp contrast, samples of *C. zenithicus* showed a high degree of overlap with the other forms, especially with *C. artedi* and *C. hoyi* (Figure S3).

**Table 3.** Percentage of overlap in isotopic niches between pairings of *C. artedi*, *C. hoyi*, *C. kiyi* and *C. zenithicus* based on the overlap of corrected standard ellipse areas. The values are the percentage of overlap of a given form in columns in the isotopic niche space of the other forms in rows.

| Species | *C. artedi* | *C. hoyi* | *C. kiyi* | *C. zenithicus* |
| --- | --- | --- | --- | --- |
| *C. artedi* | - | 8.80 | 0.00 | 29.22 |
| *C. hoyi* | 8.53 | - | 24.66 | 29.23 |
| *C. kiyi* | 0.00 | 5.82 | - | 5.18 |
| *C. zenithicus* | 90.04 | 87.26 | 65.58 | - |

### Transcriptomics

Raw reads for all individuals are available in the Sequence Read Archive of NCBI, as part of BioProject PRJNA659559. After removal of low-quality reads and trimming adaptors, we obtained an average of 46 million reads per individual (SD ± 16,414,234; Table S3). The transcriptome assembly with Trinity resulted in 1,235,811 assembled contigs for *C. artedi*, 1,504,437 for *C. hoyi*, 1,703,560 for *C. kiyi* and 1,645,826 for *C. zenithicus*. After summarizing with Transdecoder and CD-hit, the final transcriptomes resulted in 167,179 assembled contigs for *C. artedi*, 167,135 for *C. hoyi*, 137,947 for *C. kiyi* and 134,387 for *C. zenithicus* (Table S4). The resulting transcriptome is available in the Transcriptome Shotgun Assembly Sequence Database of NCBI (GIUL00000000). RNAseq reads from all species (eight individuals per species, seven for *C. zenithicus*) were mapped to the transcriptome of *C. artedi* (167,135 contigs). After quality filtering with BCFtools, the resulting matrix was comprised of 22,285 high-confidence SNPs, with no missing data.

The molecular diversity indexes for all 31 samples were *H*_*o*_= 0.27, *H*_*e*_ = 0.22, *F*_*is*_ = −0.26, *N*_*a*_ = 147,841 and θ= 0.73 (Table S5). Average *F*_*st*_ across all samples was 0.018, and *Gtest* results (*p*<0.01) indicate that the null hypothesis of panmixia was rejected, suggesting the presence of genetic differentiation among species. Pairwise *F*_*st*_ estimates showed significant, albeit low, levels of genetic differentiation between species (Table 4). The largest differences were found between *C. kiyi* and *C. zenithicus* (*F*_*st*_= 0.025, *p*< 0.01, Figure 4S) and *C. artedi* vs. *C. kiyi* (*F*_*st*_ = 0.024, *p*< 0.01; Figure 3a). The lowest level of genetic divergence was found between *C. hoyi* and *C. zenithicus* (*F*_*st*_= 0.00, *p>0.788*; Figure 3b).

**Table 4.** Estimates of pairwise genetic differentiation (Weir-Cockerham *F*_*st*_) among *C. artedi, C. hoyi, C. kiyi* and *C. zenithicus* of Lake Superior (below diagonal) and corresponding p-values (above the diagonal).

| Species | *C. artedi* | *C. hoyi* | *C. kiyi* | *C. zenithicus* |
| --- | --- | --- | --- | --- |
| *C. artedi* | - | <0.001 | <0.001 | <0.001 |
| *C. hoyi* | 0.015 | - | <0.001 | 0.788 |
| *C. kiyi* | 0.024 | 0.023 | - | <0.001 |
| *C. zenithicus* | 0.017 | 0.000 | 0.025 | - |

**Figure 3.**
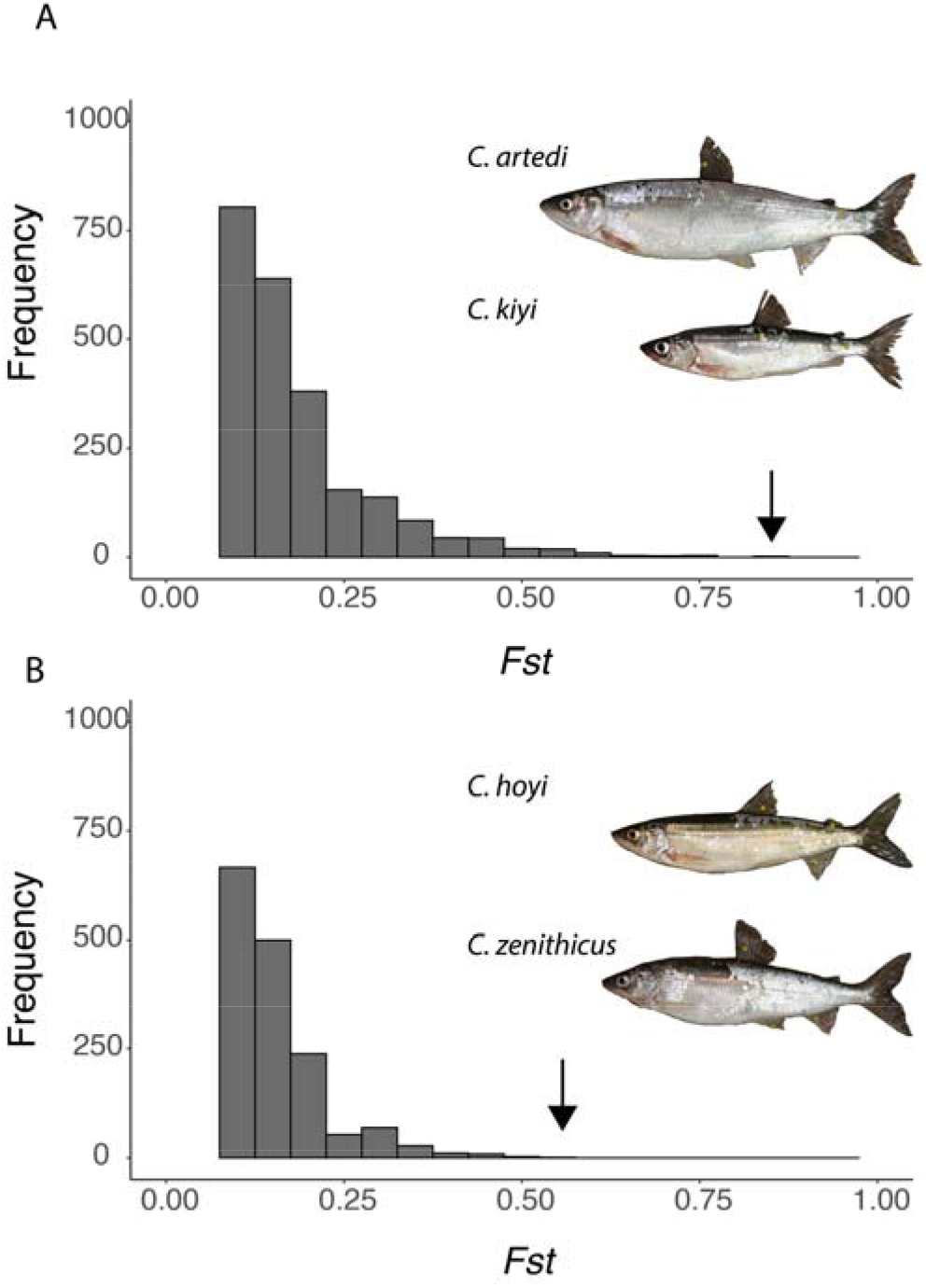
Histograms of pairwise *F*_*st*_ for individual genes in pairwise comparisons: between (A) the shallow-water *C. artedi* and deep-water *C. kiyi*, and (B) the two species with the lowest levels of divergence *C. zenithicus* and *C. hoyi*. The X-axis represents values of *F*_*st*_ (with binning of 0.05) and arrows represent the gene with the largest *F*_*st*_ (0.88 for *C. artedi* and deep-water *C. kiyi*; 0.54 for *C. zenithicus* and *C. hoyi*). The Y-axis represents frequency of individuals in log-scale. Photographs by D.L. Yule.

**Figure 4.**
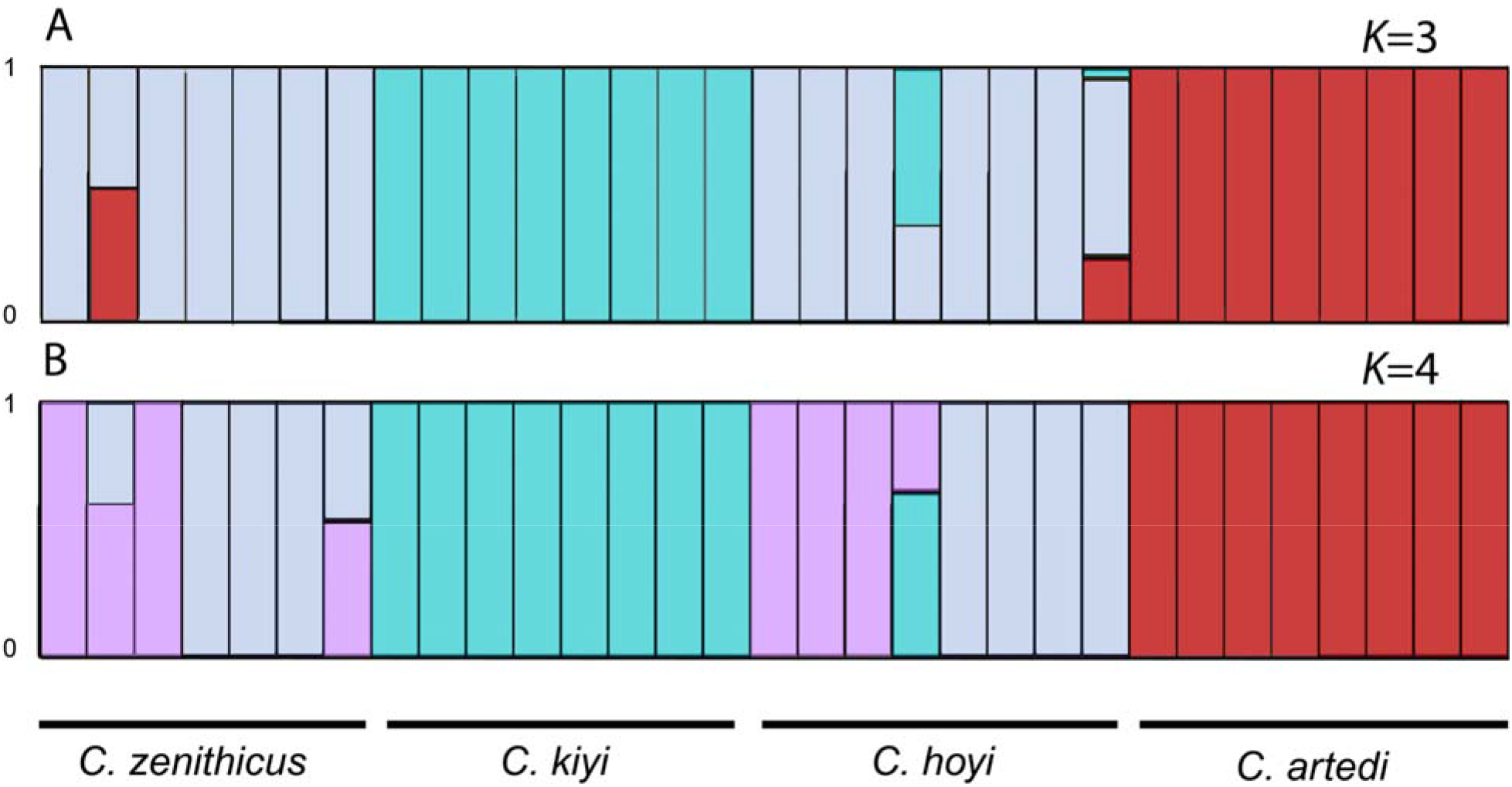
Admixture plots for *Coregonus artedi*, *C. hoyi*, *C. kiyi* and *C. zenithicus* using 22,285 high-confidence SNPs, simulating (A) three groups (*K*=3) and (B) four groups (*K*=4).

The smaller panel of SNPs derived from orthologous sequences from Orthofinder (7,898 SNPs) resulted in *F*_*st*_ values comparable to those estimated with the full dataset (Table S7). This confirms our initial observations of differentiation between the closely related groups, suggesting the significant *F*_*st*_ values are not simply derived from paralogous sequences. This is a relevant distinction to make in salmonids, given their duplicated genome.

The PCA showed concordant patterns of variation with morphology and isotopes, but with greater separation among species (Figure 2c). The largest axis of differentiation (PC1=5.44% of variance explained) was between samples of *C. kiyi* versus the rest of the groups (Figure 2c). Meanwhile PC2 explained the differences between *C. artedi* with the rest of the groups (PC2=4.58%). The SNP data showed complete separation between *C. artedi* and *C. zenithicus*, which was not observed for isotopes and morphology. As with other analyses, the PCA showed overlap between samples of *C. hoyi* and *C. zenithicus.* The PCA also showed overlap for individuals of *C. hoyi* and *C. kiyi*, and between *C. artedi* and *C. zenithicus* (Figure 2c). Meanwhile, no overlap was observed between *C. kiyi* and *C. artedi*.

The population assignment test was consistent with the presence of three groups, with clear separation among *C. artedi*, *C. kiyi*, and a cluster of *C. hoyi* and *C. zenithicus* (Figure 4a), consistent with the PCA results. When we repeated the analysis with *K*= 4, the groups formed by individuals of *C. artedi* and *C. kiyi* remained cohesive (as with *K=3*), yet the samples of *C. hoyi* and *C. zenithicus* are split into two mixed groups (Figure 4b). For the estimates of *K*=3, evidence of admixture was observed between *C. hoyi* and *C. kiyi*, *C. artedi* and *C. zenithicus*, and *C. artedi* and *C. hoyi* (Figure 4a).

The three-population analysis revealed the group formed by *C. hoyi* and *C. zenithicus* is admixed, as the Z-scores ranged from −9.72 to −14.79 (Table S6). Even when we detected potential hybrids between other species with the PCA and assignment test (i.e., *C. hoyi* and *C. kiyi*, *C. artedi* and *C. zenithicus*), these comparisons showed no significant hybridization with the *f3 statistic*. This is probably because admixture was only observed for one individual of the compared groups (Table S6).

The pairwise *Fst* comparisons between species also allowed detecting highly differentiated genes between species (Supplementary Data 1-6). Thus, the most differentiated genes for each comparison were: *Fork-head Box Q1a* (*Fst*=0.74) between *C. artedi* and *C. hoyi* (Figure S4); *Rhodopsin* (*Fst*=0.83) between *C. artedi* and *C. kiyi* (Figure 3a); *RCC1 Domain Containing 1* (*Fst*=0.64) for *C. artedi* and *C. zenithicus* (Figure S4); *Zinc Finger MYM-Type Containing 2* (*Fst*=0.86) for *C. hoyi* and *C. kiyi* (Figure S4); *Spectrin Beta, Non-Erythrocytic 2* (*Fst*=0.54) for *C. hoyi* and *C. zenithicus* (Figure 3b); *Vacuolar Protein Sorting 13 Homolog A* (*Fst*= 0.78) for *C. kiyi* and *C. zenithicus* (Figure S4). The analysis of selection with Ohana revealed 68 genes with a Log-likelihood ratios> 10 (“divergent selection”) and 12 loci with Log-likelihood ratios> 15 (“very high divergent selection”; Cheng et al., 2016; Supplementary Data 7). These could be associated with ecological and morphological functions, such as regulation of epithelial formation (*Desmoplakin; Keratin Type 1*), cell morphology and organization (*Zinc Finger MYM-type 2; Peroxiredoxin-1*), development of nervous system (*Glutathione-specific gamma-glutamylcyclotransferase 1; cAMP-responsive element modulator*), metabolism of carbohydrates (*Protein phosphatase 1 regulatory subunit 3C*), vision (*Rhodopsin*), and cellular organization during development (*Fork-head box protein F1*). In addition, we explored if there was overlap between the analysis of Ohana (68 genes) and the top 50 differentiated genes for each of the pairwise *Fst* comparisons (i.e. 6 pairwise comparisons; 300 genes in total). This comparison resulted in 47 genes that showed both high *Fst* and potential divergent selection across all ciscoes (Supplementary Data 8).

GO enrichment analyses with the Mann-Whitney Test for the pairwise *Fst* estimates showed 17 significant GO terms (FDR<10%) across four different pairwise comparisons (Supplementary Data 9). The comparison of *C. artedi* vs. *C. zenithicus* showed the most enrichment, with seven terms for Biological Process, two for cellular component, and five for Molecular Function. This included the terms membrane organization (Biological Process), and secondary active transmembrane transporter activity (Molecular Function). Other comparisons with significant results (FDR<10%) for only one category were: *C. artedi vs. C. hoyi* (Molecular Function: Rho GTPase binding), *C. hoyi* vs. *C. zenithicus* (Biological Process: cell chemotaxis, plasma membrane raft assembly); *C. kiyi* vs. *C. zenithicus* (Cellular Component: plasma membrane region, phosphatidylinositol 3-kinase complex).

## DISCUSSION

Understanding the dynamics of speciation can be enhanced by studying taxa with differences in phenotypic and ecological traits, but modest genetic differentiation. This is the case in Great Lakes coregonines, which are characterized by having complex evolutionary origins, as they show variation in morphological traits due to plasticity, selection, and/or hybridization. For taxonomically-complex taxa such as coregonines, integrating new results with previously published studies is hindered by a long history of researchers either implicitly or explicitly applying different taxonomic frameworks, often in the absence of museum voucher specimens. Thus, it is often not possible to reconcile whether apparent differences across space, time, or taxa, are due to true biological differences or reflect the taxonomic identifications employed. In taxa where this complexity occurs, we suggest that researchers either (1) explicitly describe the methods and morphological characters used in assigning individuals to taxa, (2) include linked morphological/ecological, and genetic data from the same specimens, or (3) deposit voucher specimens in natural history museums to serve as an historical record (e.g., see Schmidt et al. 2009). With this in mind, the present study applied an integrative approach to characterize morphological, ecological, and genetic analyses to the same set of individuals to disentangle the differences found within this economically and ecologically relevant group of fishes. The linked genome-to-phenome approach applied in this study will allow for greater comparability across studies and more robust analyses of variation in Great Lakes ciscoes. Limitations of this approach include the necessarily small sample sizes, but with DNA sequencing costs diminishing over time, this issue should eventually be overcome. Overall, understanding the relationships within *Coregonus* is highly relevant today, as overfishing, habitat degradation and invasive species have led to declines in abundance and diversity for decades (Eshenroder et al., 2016).

### Isotopic measures show agreement with previous studies

Based on isotopic measurements, it appears that the main driving mechanism of dietary partitions is depth preference, which in turn influences prey availability. These findings confirm previous observations that muscle derived δ^13^C concentrations in *C. kiyi* and *C. hoyi*/*C. zenithicus* are less depleted than in *C. artedi*. This reflects preference for pelagic habitat and primary consumption of limnetic zooplankton during summer for the latter (Sierszen et al., 2014; Rosinski et al., 2020). This result is expected given the near-obligate pelagic existence of adult individuals of *C. artedi* during summer stratification (Rosinski, Vinson, & Yule, 2020). Meanwhile, high average δ^13^C and δ^15^N values of *C. kiyi* are consistent with a diet dominated by *Mysis diluviana* (Gamble, Hrabik, Stockwell, & Yule, 2011a; Ahrenstorff et al., 2011; Sierszen et al., 2014) and preference for deep waters (Rosinski et al., 2020). The intermediate position of *C. hoyi* suggest they occupy intermediate depths and are opportunistic feeders with a diet that is obtained from pelagic and benthic zones. This result is also consistent with previous studies that showed *C. hoyi* change their diet seasonally (*Diporeia* in spring, *Mysis diluviana* in summer, and *Daphnia* spp. and calanoid copepods in autumn; Gamble, Hrabik, Stockwell, & Yule, 2011b). When compared to the other species, the isotopic values of *C. zenithicus* (Figure 2b) suggest that some individuals in this group may be obtaining part of their nutrition from the pelagic zone, through consumption of zooplankton. Samples of *C. zenithicus* from this study had terminal mouths similar to *C. artedi*, which could explain the wide breath of dietary components compared to the other groups. Both the present study and Schmidt et al. (2009) have *C. zenithicus* and *C. hoyi* occupying similar trophic positions, and both show *C. zenithicus* obtaining nutrition from more δ^13^C depleted prey resources relative to *C. hoyi.* Yet, Schmidt et al. (2009) showed these two species occupied unique isotopic space (especially for their Lake Superior samples of 1972-1998), while we obtained far greater overlap. Given these contrasting results, it is necessary to expand the studies on the dietary overlap between *C. hoyi* and *C. zenithicus* among different populations of Lake Superior.

### Multiple lines of evidence support divergence of *C. artedi* and *C. kiyi*

The forms that showed the highest levels of divergence across all measured traits were the shallow water *C. artedi* and the deep-water *C. kiyi.* One of the most striking morphological differences between the two is the size of the eye, which is much larger in *C. kiyi* relative to the rest of their bodies (Table 1). Genetic differences were also associated with vision, as two highly divergent genes between the two species were *Rhodopsin* (RHO; *F*_*st*_=0.86) and *Homeodomain-interacting protein kinase 2* (HIPK2; *F*_*st*_=0.55). *Rhodopsin* is a key photoreceptor in low light conditions, and differences in this gene have been previously reported for the two species (Eaton et al., 2020), as well as for other lineages exposed to dim-lit environments (Hill et al., 2019; Musilova et al., 2019; Schott, Refvik, Hauser, López-Fernández, & Chang, 2014). *Homeodomain-interacting protein kinase 2* is associated with tissue growth during development, including eye size and lens formation in mice (Inoue et al., 2010; Poon, Zhang, Lin, Manning, & Harvey, 2012). Additional loci that were highly differentiated between *C. artedi* and *C. kiyi* include: *Insulin Growth Factor 1 Receptor* (IGRF1; *F*_*st*_=0.87) which is essential for embryonic growth and cell proliferation of vertebrates (Schlueter, Peng, Westerfield, & Duan, 2007); *Plasminogen Activator* (PLAT; *F*_*st*_=0.73), associated with tissue formation and modification in fishes (Bader, et al. 2012); *Dystroglycan* (DAG1; *F*_*st*_=0.62) which has many functions including muscle development in zebrafish (Parsons, Campos, Hirst, & Stemple, 2002); and *cAMP-responsive element modulator* (CREM; *F*_*st*_=0.62) important for neurogenesis and neuro-plasticity of vertebrates (Mioduszewska, Jaworski, & Kaczmarek, 2003). These are candidate genes that could exhibit adaptive genetic variants associated with habitat preferences of the two species, and the morphological differences found between them.

The PCL/PPD and PVL/PAD ratios were also considerably smaller in *C. artedi* compared to *C. kiyi*. Previous studies suggest that this is a convergent adaptation across lacustrine fishes, where large bodies and smaller fins provide advantages for maintaining buoyancy in pelagic environments with little swimming (i.e., *C. artedi*), while small bodies with longer fins favor swimming at depth (*C. kiyi*; Eshenroder, Sideleva, & Todd, 1999). Overall, the results from our study suggest concordance between morphological, genetic, and ecological differences for a shallow-water pelagic species (*C. artedi*) and the deep-water species (*C. kiyi*). Future studies are needed to confirm if there is a relationship between the aforementioned genes and morphological and ecological differences detected among the groups.

### Lack of differentiation between *C. zenithicus* and *C. hoyi*

One of the revealing findings of our study was the limited differentiation between samples initially identified as *C. zenithicus* and *C. hoyi*. At first sight, the isotopic analysis suggests *C. hoyi* are obtaining more nutrition from the benthic zone, but this signal could be confounded by seasonal variation. Further, samples identified as *C. zenithicus* and *C. hoyi* showed no significant differences in the overall *F*_*st*_, and the *f3-statistic* indicated rampant admixture between the two taxa. Still, the pairwise comparisons of locus specific *F*_*st*_ showed differentiation for genes associated with nerve growth and neural development (*Spectrin Beta Non-Erythrocytic 2*, *F*_*st*_=0.54; *Growth Associated Protein 43*, *F*_*st*_=0.47; Benowitz & Routtenberg, 1997; Lise et al., 2012), as well as cell proliferation in vertebrates (*Tensin 2*; *Fst*=0.49; Hafizi, Ibraimi & Dahbäck, 2005). These highly differentiated loci did not influence the low overall *F*_*st*_ as they represent a very small proportion of the sampled loci. Future studies across populations of *C. hoyi* and *C. zenithicus* are necessary to understand the full extent of the genomic divergence between the two species.

The key phenotypic traits used in our study to distinguish *C. zenithicus* and *C. hoyi* were lower jaw position (*C. zenithicus*: included or terminal; *C. hoyi*: extended) and gill raker counts (*C. zenithicus* mean ± SD = 39.5 ± 2.3; *C. hoyi* = 42.4 ± 2.1; Eshenroder et al., 2016), both of which are commonly used in the field to identify these species. Previous studies also suggest that both species can be differentiated through pre-maxilla angle, which are ~40° for *C. hoyi*, and between 60°-65° for *C. zenithicus* (Eshenroder et al., 2016). The range of pre-maxillary angles for our samples was on average 35.88° (SD ±4.05) for *C. hoyi*, and 35.52° (SD ±3.14) for *C. zenithicus*. Thus, it is possible that we did not collect samples of *C. zenithicus* in our study and that all three measurements (i.e., lower jaw position, gill raker morphology, and pre-maxillary angles) should be employed when differentiating the two species.

Alternatively, it is possible that these two taxa are now forming a hybrid-swarm that is morphologically and genetically very similar in certain areas of the Great Lakes. This hypothesis stems from the recent collapse of *C. zenithicus* in Lake Superior. Koelz (1929) reported that *C. zenithicus* was the dominant deep-water cisco in Lake Superior in 1921-1922 representing 90% of the ciscoes caught in bottom set gillnets (Eshenroder et al., 2016). By 2001-2003 they represented only 4% of ciscoes caught in nearshore and offshore bottom trawl samples (Gorman & Todd 2007). Declines in *C. zenithicus* abundance, coupled with the propensity for *C. hoyi* to expand into areas previously occupied by other deep species (Bronte et al., 2010; Bunnell et al., 2012), could have promoted hybridization. In our study, we also show evidence of admixture between 1 or 2 individuals *C. hoyi* and *C. kiyi*, as well as *C. zenithicus* and *C. artedi*. Previous studies have demonstrated that hybridization is possible between closely related coregonines in both North America and Europe (Hudson, LundsgaardLHansen, Lucek, Vonlanthen, & Seehausen, 2017; Hudson, Vonlanthen, & Seehausen, 2011; Kirtiklis & Jankun, 2006; Todd & Stedman, 1989), and successful crosses of *C. artedi* and *C. hoyi* have been made in captivity (W. Stott, personal communication). Further, hybridization has been suggested as one of the drivers of morphological changes in *C. hoyi* in Lake Huron after the decline of other species in the complex (Todd & Stedman, 1989). This scenario of speciation reversal in *C. zenithicus*, where morphological diversity could have been lost due to both human-mediated stressors and admixture with closely related species, could represent an example of how commercially and ecologically relevant lineages have been lost in recent decades. It remains to be determined if other populations of *C. zenithicus* and *C. hoyi* in Lake Superior are also admixed, given the limited number of samples included in the present study. This is especially relevant considering previous surveys have reported finding phenotypically distinct *C. zenithicus* in Canadian waters of Lake Superior (Pratt & Chong, 2012; Pratt, 2012).

### Concordance among morphology, ecology and genetics

This study exemplifies how analyzing the same specimens with multiple approaches can enhance our understanding of the relationships among the *C. artedi* complex. Our results showed concordance for morphological, ecological, and genetic divergence for three of the four species studied. The highest levels of differentiation for all the analyzed traits was observed for *C. kiyi*, which is the species typically found in deeper waters of Lake Superior. This is expected given that adaptation to life at depth has been reported as one of the main drivers of differentiation for lacustrine fishes, including African cichlids (Albertson, 2008; Schliewen et al., 2001), grayling (Olson, Krabbenhoft, Hrabik, Mendsaikhan, & Jensen, 2019), sculpins of Lake Baikal (Kontula, Kirilchik, & Väinölä, 2003), lake trout in Great Lakes (Perreault-Payette et al., 2017), and European coregonines (Vonlanthen et al., 2009). In Lake Superior, our results suggest adaptation to depth has resulted in a larger eye and modifications of the visual system (e.g., *Rhodopsin*; Eaton et al. 2020), as well as a diet centered around the deep macro-invertebrate *Mysis*, when compared to its shallower counterparts.

This study also suggests that another driver of differentiation for *Coregonus* is the benthic/pelagic axis, which is also known to lead to the differentiation of lacustrine species (e.g., cichlids: Hulsey, Roberts, Loh, Rupp, & Streelman, 2013; perch: Svanbäck & Eklöv, 2004). This is evidenced by the results of stable isotope analysis, where *C. artedi* appears to have mostly a pelagic diet, while the remaining groups had a combination of pelagic/benthic elements (Sierszen et al., 2014). Observed dietary preferences could also be linked to the differences in the angle of the mouth, as having terminal mouths can be associated with feeding on pelagic zooplankton (*C. artedi*), while having a more angled mouth could provide advantages when feeding at depth (*C. kiyi* and *C. hoyi;* Etheridge, Bean, Maitland, Ballantyne, & Adams, 2012). It is important to highlight that feeding morphologies can change for this group based on competition and prey availability, as populations of *C. artedi* and *C. zenithicus* can show considerable differences in feeding structures across the Great Lakes depending on the presence/absence of congeners (Turgeon et al., 2016).

Overall, the results of our study suggest that differentiation across *Coregonus* could be promoted by habitat preferences (i.e., depth and benthic/pelagic habit) and dietary partitions, both of which are well known drivers of diversification of multiple fish groups (Bernardi, 2013).

### Status of the Great Lakes and conservation of genetic resources

Extensive efforts are being made to improve the conditions for organisms that inhabit the Great Lakes, including the reduction of contaminants (e.g., PCBs) and controlling densities of invasive species (e.g., Siefkes et al. 2013 and *https://www.epa.gov/greatlakes/lakewide-action-and-management-plans-great-lakes*) and reestablishment of native fishes (e.g., Muir et al. 2012). Along with these efforts, it is essential for managers to identify and monitor functional genomic variants associated with ecological and morphological differences of coregonines. This is critical as it allows for the maintenance of the evolutionary potential of the *C. artedi* complex across the Great Lakes. Further, evaluating individual genes and molecular pathways associated with ecological divergence across the Great Lakes will allow us to understand if some populations are at higher risk of speciation reversal due to population declines or changing environmental conditions. These approaches should be implemented across lakes that have different faunal compositions, and especially in lakes that have fewer sympatric species of *Coregonus* than Lake Superior (e.g., Lakes Michigan and Huron), as both of these factors are known to influence the amount of divergence among coregonines (e.g., Turgeon et al., 2016).

## Conclusions

In this study, we analyzed patterns of divergence among four species of *Coregonus* from Lake Superior and found concordance among morphological, ecological and genomic approaches. Based on the morphological analyses, we confirmed previous observations that eye size, fin length and length of the lower jaw explain most of the variation between species. There are clear contrasts in the dietary preferences of *C. artedi* compared to *C. kiyi*, while observed differences between *C. hoyi* and *C. zenithicus* appear to be largely influenced by seasonality. Transcriptomic analyses were able to distinguish between three of the four species, and high levels of differentiation in genes associated with body shape, eye size, fin shape, vision, organization of the nervous system and early development. There was evidence of hybridization between samples morphologically identified as *C. hoyi* and *C. zenithicus* and future studies should assess the extent of this admixture in Lake Superior. Overall, the integrative approach applied in this study exemplifies how multiple lines of evidence can help elucidate the relationships among sympatric groups with complex evolutionary history, such Great Lakes *Coregonus* species.

## Acknowledgements

We thank the U.S. Geological Survey Research Vessel Kiyi Captain Joe Walters, First Mate Keith Peterson, and Engineer Charles Carrier. Special thanks to Ryan Menenbroeker who measured our specimens, Jean Adams for analyzing the morphometric data, and to Caroline Rosinski, Mark Vinson, and Andrew Muir for providing helpful insight. We would like to thank Tianying Lan, Nathan Backenstose, Katherine Eaton, and Jessie Pelosi for their help with the genetic analyses and data handling. Early drafts of this manuscript were improved with solicited reviews by Tom Pratt and Gary Longo. Any use of trade, product, or firm names is for descriptive purposes only and does not imply endorsement by the U.S. Government. Funding was provided by start-up funds to TED from Wayne State University and the Great Lakes Fishery Commission (Award no. 2018-KRA-44073).

## Data Accessibility

The raw reads for all individuals and the assembled transcriptome of *Coregonus artedi* are available in NCBI as BioProject PRJNA659559. The bioinformatic scripts used for the transcriptomic analyses are available at github.com/evofish.

## Author Contributions

TED and TJK conceived the study; DY and LE collected the samples; DLY, LE, and TJK collected the data; MAB, DY, WS and TJK analyzed the data and interpreted the results; the manuscript was written by MAB and DY and all the co-authors contributed to the final version.

