## Supplement for "Transcriptomic divergence predicts morphological and ecological variation underlying an adaptive radiation"

**SUPPLEMENTAL TABLES**

**Table S1**. Length, weight, sex and collection sites for samples included in this study. Site number corresponds to those on Figure 1. CA= *Coregonus artedi*; CH= *C. hoyi*; CK= *C. kiyi*; CZ= *C. zenithicus*.

| **Sample** | **Site** | **Species** | **Date** | **Length** | **Weight** | **Sex** | **Latitude** | **Longitude** |
| --- | --- | --- | --- | --- | --- | --- | --- | --- |
| **CA01** | 1 | *C. artedi* | 8/25/15 | 325 | 264 | N/A | 46.80628333 | -90.78238333 |
| **CA02** | 5 | *C. artedi* | 8/29/15 | 340 | 296 | N/A | 46.95410833 | -90.47148333 |
| **CA03** | 5 | *C. artedi* | 8/29/15 | 326 | 272 | N/A | 46.95410833 | -90.47148333 |
| **CA04** | 5 | *C. artedi* | 8/29/15 | 300 | 216 | N/A | 46.95410833 | -90.47148333 |
| **CA05** | 1 | *C. artedi* | 8/25/15 | 324 | 276 | N/A | 46.80628333 | -90.78238333 |
| **CA06** | 15 | *C. artedi* | 11/3/15 | 372 | 365 | Male | 48.46105800 | -88.89977500 |
| **CA07** | 15 | *C. artedi* | 11/3/15 | 324 | 257 | Male | 48.46105800 | -88.89977500 |
| **CA08** | 14 | *C. artedi* | 11/4/15 | 341 | 322 | Male | 48.43200833 | -89.01235000 |
| **CH01** | 4 | *C. hoyi* | 5/19/15 | 250 | 109 | Female | 46.89205417 | -90.53255833 |
| **CH02** | 3 | *C. hoyi* | 5/29/15 | 241 | 97 | Female | 46.88540833 | -91.21529167 |
| **CH03** | 3 | *C. hoyi* | 5/29/15 | 267 | 124 | Female | 46.88540833 | -91.21529167 |
| **CH04** | 3 | *C. hoyi* | 5/29/15 | 222 | 78 | Female | 46.885401833 | -91.21529167 |
| **CH05** | 4 | *C. hoyi* | 5/19/15 | 200 | 56 | Male | 46.89205417 | -90.53255833 |
| **CH06** | 7 | *C. hoyi* | 5/20/15 | 226 | 75 | Female | 46.97675833 | -90.45365833 |
| **CH07** | 3 | *C. hoyi* | 5/29/15 | 270 | 125 | Female | 46.88540833 | -91.21529167 |
| **CH08** | 3 | *C. hoyi* | 5/29/15 | 267 | 122 | Female | 46.88540833 | -91.21529167 |
| **CK01** | 4 | *C. kiyi* | 5/19/15 | 238 | 82 | Female | 46.89205417 | -90.53255833 |
| **CK02** | 12 | *C. kiyi* | 7/8/15 | 240 | 99 | N/A | 47.49667500 | -89.99912500 |
| **CK03** | 10 | *C. kiyi* | 7/8/15 | 241 | 106 | N/A | 47.15711667 | -89.96870833 |
| **CK04** | 11 | *C. kiyi* | 7/10/15 | 227 | 72 | N/A | 47.41659167 | -88.46418333 |
| **CK05** | 13 | *C. kiyi* | 7/7/15 | 221 | 65 | N/A | 47.53425000 | -90.54291667 |
| **CK06** | 13 | *C. kiyi* | 7/7/15 | 228 | 74 | N/A | 47.53425000 | -90.54291667 |
| **CK07** | 12 | *C. kiyi* | 7/8/15 | 206 | 50 | N/A | 47.49667500 | -89.99912500 |
| **CK08** | 10 | *C. kiyi* | 7/8/15 | 199 | 38 | N/A | 47.15711667 | -89.96870833 |
| **CZ01** | 2 | *C. zenithicus* | 5/28/15 | 267 | 135 | Female | 46.81871667 | -91.41867500 |
| **CZ02** | 9 | *C. zenithicus* | 8/30/15 | 251 | 110 | N/A | 47.03325800 | -90.60138300 |
| **CZ03** | 9 | *C. zenithicus* | 8/30/15 | 272 | 145 | N/A | 47.03325800 | -90.60138300 |
| **CZ04** | 6 | *C. zenithicus* | 8/30/15 | 256 | 134 | N/A | 46.96980833 | -90.69405833 |
| **CZ05** | 2 | *C. zenithicus* | 5/28/15 | 255 | 108 | Female | 46.81871667 | -91.41867500 |
| **CZ06** | 3 | *C. zenithicus* | 5/29/15 | 230 | 78 | Female | 46.88540833 | -91.21529167 |
| **CZ07** | 8 | *C. zenithicus* | 8/30/15 | 296 | 172 | N/A | 47.03245000 | -90.60183300 |
| ***CZ08**** | 15 | *C. zenithicus* | 11/3/15 | 228 | 94 | Female | 48.46105800 | -88.89977500 |

**CZ08 was not included in the final analyses, as it was misidentified in the field*.

**Table S2.** Full description of morphometric characters measured for assessment of phenotypic differences.

| Abbreviation | Measure | Description |
| --- | --- | --- |
| BDD | Body depth | Vertical distance from the origin of the dorsal fin to the ventral surface of the body. |
| DOH | Dorsal fin height | Distance from the origin of the dorsal fin to the tip of the longest ray. |
| HLL | Head length | Distance from the tip of the snout to the most extreme posterior margin of the operculum, not counting the opercular membrane, as measured parallel to the longitudinal axis of the fish. |
| MXL | Maxillary length | Distance from the most-anterior point of the premaxillary to the posterior end of the maxillary bone. |
| OOL | Orbital length (eye) | Distance between the anterior and posterior fleshy margins of the orbit with calipers anchored against margins of orbital rim |
| PAD | Pelvic-anal distance | Distance between the anterior insertions of the pelvic and anal fins. |
| PCL | Pectoral fin length | Distance from the origin of the fin to the tip of the longest ray. |
| PMA | Premaxillary angle (degrees) | Angle between the horizontal axis of the head and the symphysis of the premaxillaries. |
| POL | Preorbital length (snout) | Tip of the snout to the anterior fleshy margin of the orbital rim with calipers anchored against the rim. |
| PVL | Pelvic fin length | Distance from the origin of the fin to the tip of the longest ray. |
| PPD | Pectoral-pelvic distance | Distance between the anterior insertions of the pectoral and pelvic fins. |
| STL | Standard length | Distance from the tip of the premaxillary to the caudal flexure, i.e., the crease created when the tail is flexed. |

**Table S3.** Number of reads per individual, after removal of adaptors and low-quality reads using Trimmomatic. The average per sample was 45,819,960.44 reads (SD± 16,414,234.38). CA= *Coregonus artedi*; CH= *C. hoyi*; CK= *C. kiyi*; CZ= *C. zenithicus*.

| Sample | Reads | Sample | Reads | Sample | Reads | Sample | Reads |
| --- | --- | --- | --- | --- | --- | --- | --- |
| CA01 | 58,364,870 | **CH01** | 49,076,048 | **CK01** | 70,859,120 | **CZ01** | 53,762,858 |
| CA02 | 15,876,840 | **CH02** | 52,279,490 | **CK02** | 34,185,878 | **CZ02** | 31,640,336 |
| CA03 | 23,472,286 | **CH03** | 35,219,078 | **CK03** | 43,044,280 | **CZ03** | 50,327,228 |
| CA04 | 40,144,912 | **CH04** | 40,634,580 | **CK04** | 30,279,646 | **CZ04** | 41,406,800 |
| CA05 | 46,861,146 | **CH05** | 53,478,582 | **CK05** | 104,890,778 | **CZ05** | 57,159,472 |
| CA06 | 26,570,200 | **CH06** | 33,754,678 | **CK06** | 50,772,502 | **CZ06** | 44,180,994 |
| CA07 | 52,177,116 | **CH07** | 45,750,450 | **CK07** | 56,896,564 | **CZ07** | 56,495,418 |
| CA08 | 35,230,764 | **CH08** | 61,901,930 | **CK08** | 26,209,828 | **CZ08** | 43,334,062 |

**Table S4.** Number of contigs generated after the assembly with Trinity for each of the forms, and remaining contigs after filtering with Transdecoder and CD-Hit.

|  | Trinity Transcriptome | Transdecoder and CD-Hit Filtering |
| --- | --- | --- |
| *C. artedi* | 1,235,811 | 167,179 |
| *C. hoyi* | 1,504,437 | 167,135 |
| *C. kiyi* | 1,703,560 | 137,947 |
| *C. zenithicus* | 1,645,826 | 134,387 |
| Average | 1,522,409 | 151,662 |

**Table S5.** Molecular diversity indexes for *Coregonus artedi, C. hoyi, C. kiyi* and *C. zenithicus*: Expected Heterozygosity (*He*), Observed Heterozygosity (*Ho*), Inbreeding Coefficient (*F_is_*), Number of Alleles (*N_A_*) and Genetic Diversity (***θ***).

|  | Samples | He | Ho | F_is_ | N_A_ | *θ* |
| --- | --- | --- | --- | --- | --- | --- |
| *C. artedi* | 8 | 0.2057 | 0.2572 | -0.1885 | 36787 | 0.6317 |
| *C. hoyi* | 8 | 0.2188 | 0.2754 | -0.1986 | 37463 | 0.6434 |
| *C. kiyi* | 8 | 0.2059 | 0.2639 | -0.2209 | 36529 | 0.6172 |
| *C. zenithicus* | 7 | 0.2209 | 0.2815 | -0.2016 | 37062 | 0.6268 |
| Total | 31 | 0.2245 | 0.2695 | -0.26 | 147841 | 0.7271 |

**Table S6.** Result of the three-population test as applied by the software TreeMix V1.13 for species of the genus *Coregonus*, showing the potential for admixture of Group A, based on the genetic composition of Groups B and C. If the Z-score for Group A is negative, it is estimated that the population is the result of recent admixture from the other two populations. In this case, hybridization was only detected for *C. hoyi* and *C. zenithicus*.

| Group A | Group B | Group C | f3 statistic | Standard Error f3 | Z-Score |
| --- | --- | --- | --- | --- | --- |
| *C. hoyi* | ***C. kiyi*** | ***C. zenithicus*** | -0.0021 | 0.0002 | -12.55 |
| *C. kiyi* | ***C. hoyi*** | ***C. zenithicus*** | 0.0047 | 0.0003 | 15.79 |
| *C. zenithicus* | ***C. kiyi*** | ***C. hoyi*** | -0.0020 | 0.0002 | -9.72 |
| *C. artedi* | ***C. kiyi*** | ***C. hoyi*** | 0.0005 | 0.0002 | 2.18 |
| *C. hoyi* | ***C. kiyi*** | ***C. artedi*** | 0.0003 | 0.0002 | 1.49 |
| *C. kiyi* | ***C. hoyi*** | ***C. artedi*** | 0.0023 | 0.0003 | 8.29 |
| *C. artedi* | ***C. kiyi*** | ***C. zenithicus*** | 0.0004 | 0.0002 | 1.59 |
| *C. kiyi* | ***C. zenithicus*** | ***C. artedi*** | 0.0024 | 0.0003 | 8.80 |
| *C. zenithicus* | ***C. kiyi*** | ***C. artedi*** | 0.0003 | 0.0002 | 1.40 |
| *C. artedi* | ***C. hoyi*** | ***C. zenithicus*** | 0.0028 | 0.0002 | 15.32 |
| *C. hoyi* | ***C. zenithicus*** | ***C. artedi*** | -0.0019 | 0.0001 | -14.79 |
| *C. zenithicus* | ***C. hoyi*** | ***C. artedi*** | -0.0021 | 0.0002 | -11.61 |

**Table S7.** Estimates of pairwise genetic differentiation (Weir-Cockerham *Fst*) between *Coregonus artedi, C. hoyi, C. kiyi* and *C. zenithicus* of Lake Superior (below the diagonal) and their corresponding *p-values* (above the diagonal), using only the orthologous genes detected with Orthofinder between the four groups of interest and *C. lavaretus* (7,898 SNPs).

|  | *C. artedi* | *C. hoyi* | *C. kiyi* | *C. zenithicus* |
| --- | --- | --- | --- | --- |
| *C. artedi* | - |  |  |  |
| *C. hoyi* | 0.014 | - |  |  |
| *C. kiyi* | 0.023 | 0.019 | - |  |
| *C. zenithicus* | 0.016 | 0.000 | 0.021 | - |

**SUPPLEMENTAL FIGURES**

**Figure S1**. BUSCO assessment of transcriptome completeness for two versions of the transcriptome of *Coregonus artedi,* directly estimated from Trinity and after filtering with Transdecoder. The transcriptome of *C. artedi* was used as reference for the SNP validation and downstream comparisons for all forms of *Coregonus* spp*.*

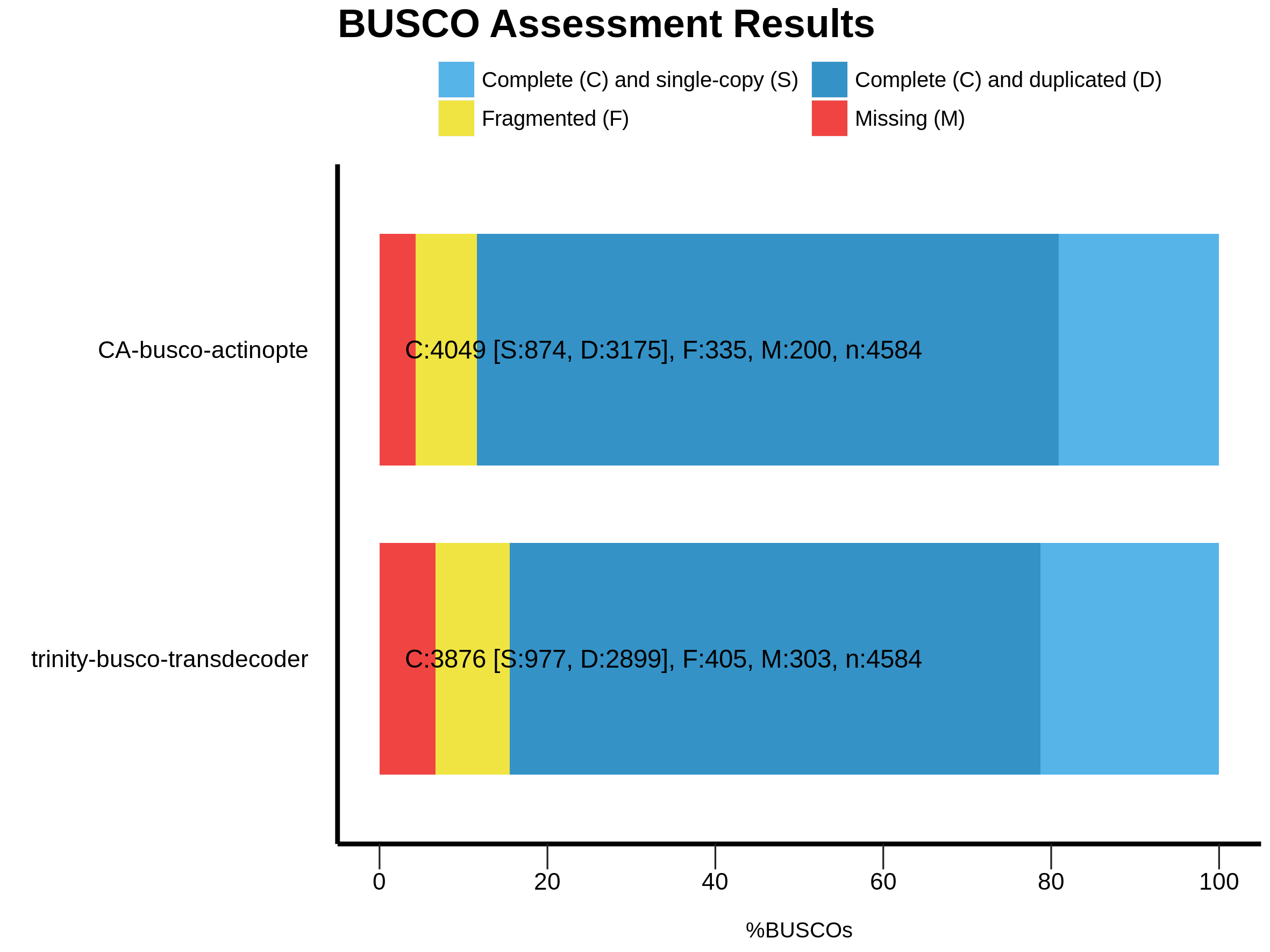

CA- Trinity and Transdecoder

CA- Trinity

**Figure S2**. Photograph (A) and Principal Component Analysis of the SNP data (B) showing the misidentified individual of *Coregonus zenithicus* (red circle). This individual showed traits that are more similar to *C. kiyi* (notably the large eyes), and the genomic analyses with SNPs confirm the identification as the latter. This individual was excluded from the final *F_st_* estimates as we had clear evidence it was a case of misidentification during field collections. The individual represented by the blue dot in the PCA was morphologically identical to other samples of *C. hoyi*, and appears to be a hybrid between the latter and *C. kiyi*. *Photograph by DL Yule*.

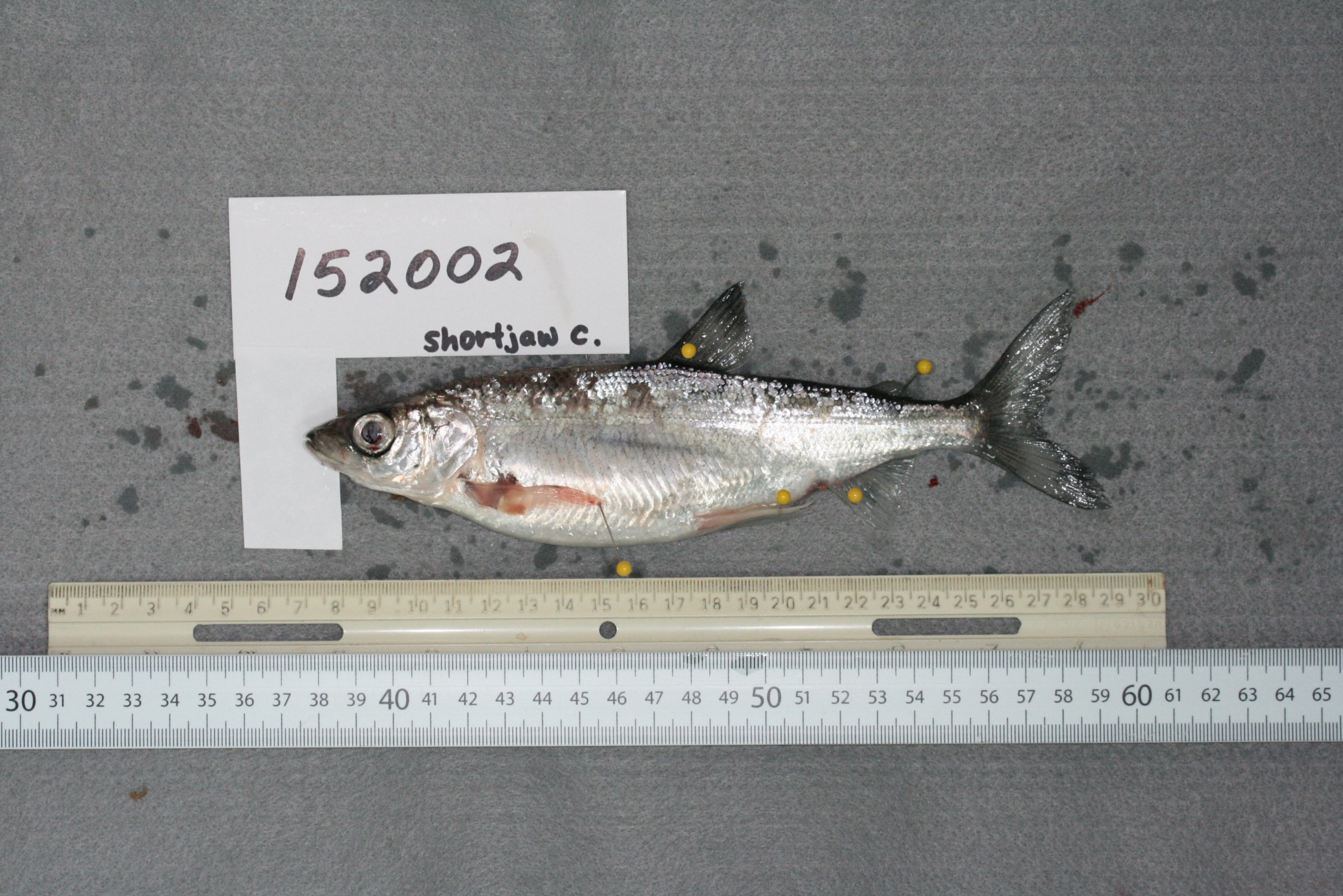

A

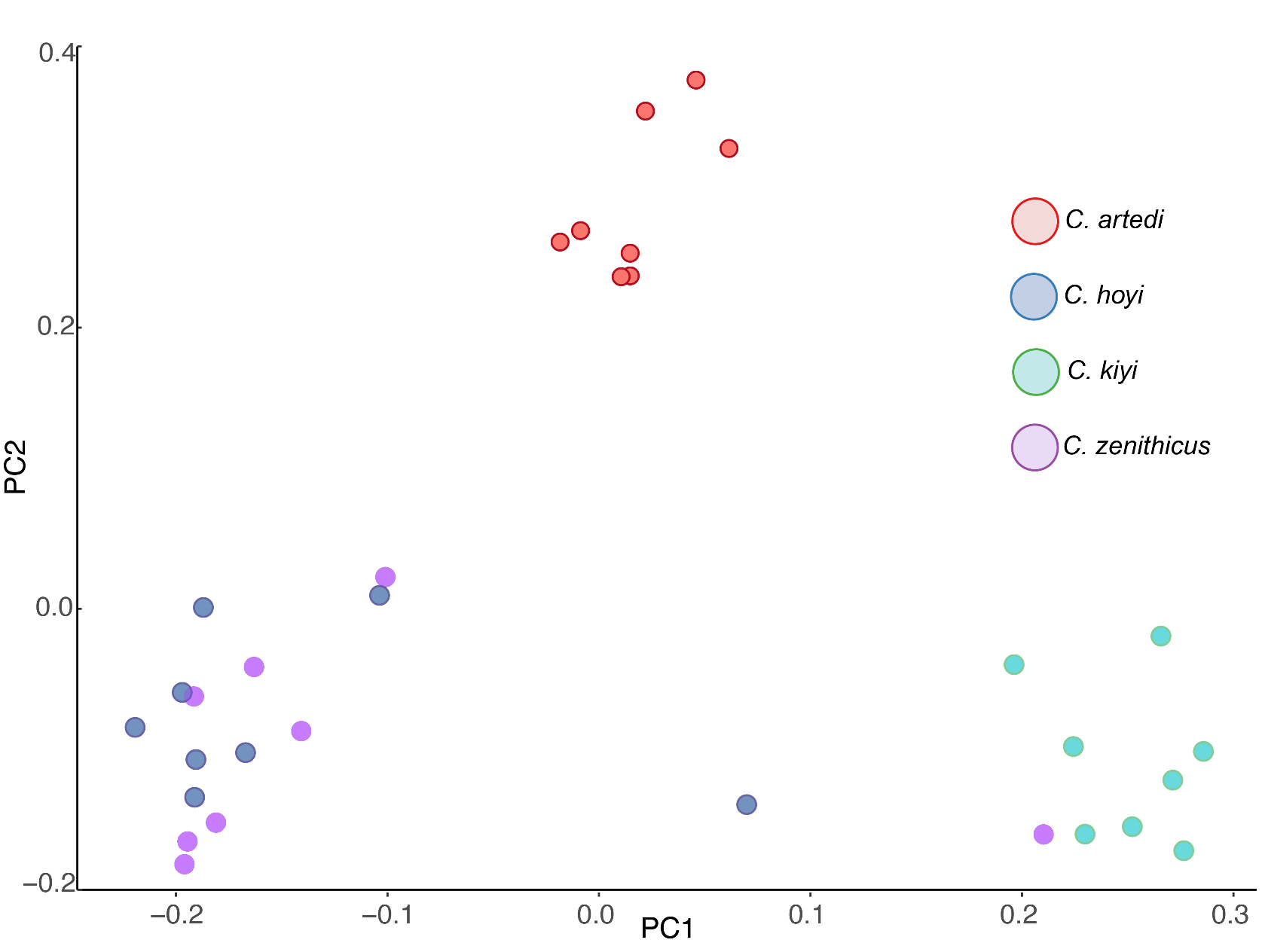
B

**Figure S3.** The δ13C_lipid free_ and δ15N bivariate plots for *C. hoyi*, *C. kiyi*, *C. artedi*, and *C. zenithicus* collected during May through November 2015 from western Lake Superior. The ellipses, called corrected Standard Ellipse Areas (SEAc) represent areas that a subsequently sampled datum would have a 95% probability of being encompassed.

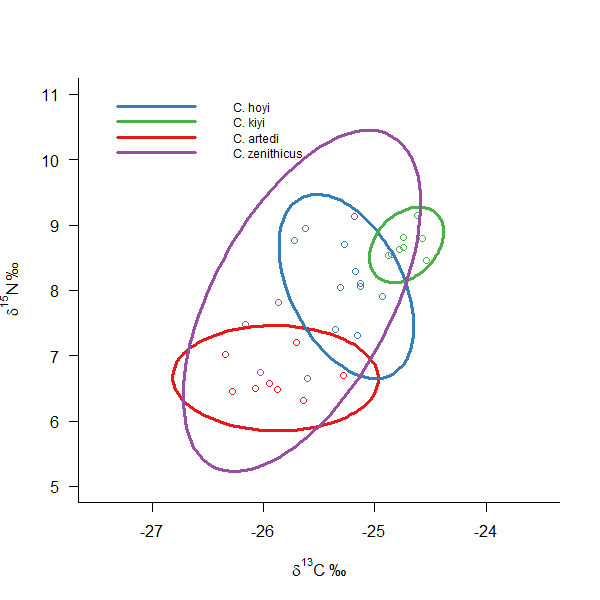

­­­

**δ^13^C_lipid free_**

**δ^13^N**

**Figure S4**. Histograms of pairwise *Fst* between: (A) *C. artedi* vs *C. hoyi* (top 1% *Fst* >0.29); (B) *C. artedi* vs *C. zenithicus* (top 1% *Fst* > 0.28); (C) *C. kiyi* vs *C. hoyi* (top *Fst* > 0.34); and (D) *C. kiyi* vs *C. zenithicus* (top *Fst* > 0.31)*.* The X-axis represents Fst estimates (binning 0.05), and arrows represent the location of the locus with the highest divergence in each comparison. Y-axis represents the number of loci in log-scale.

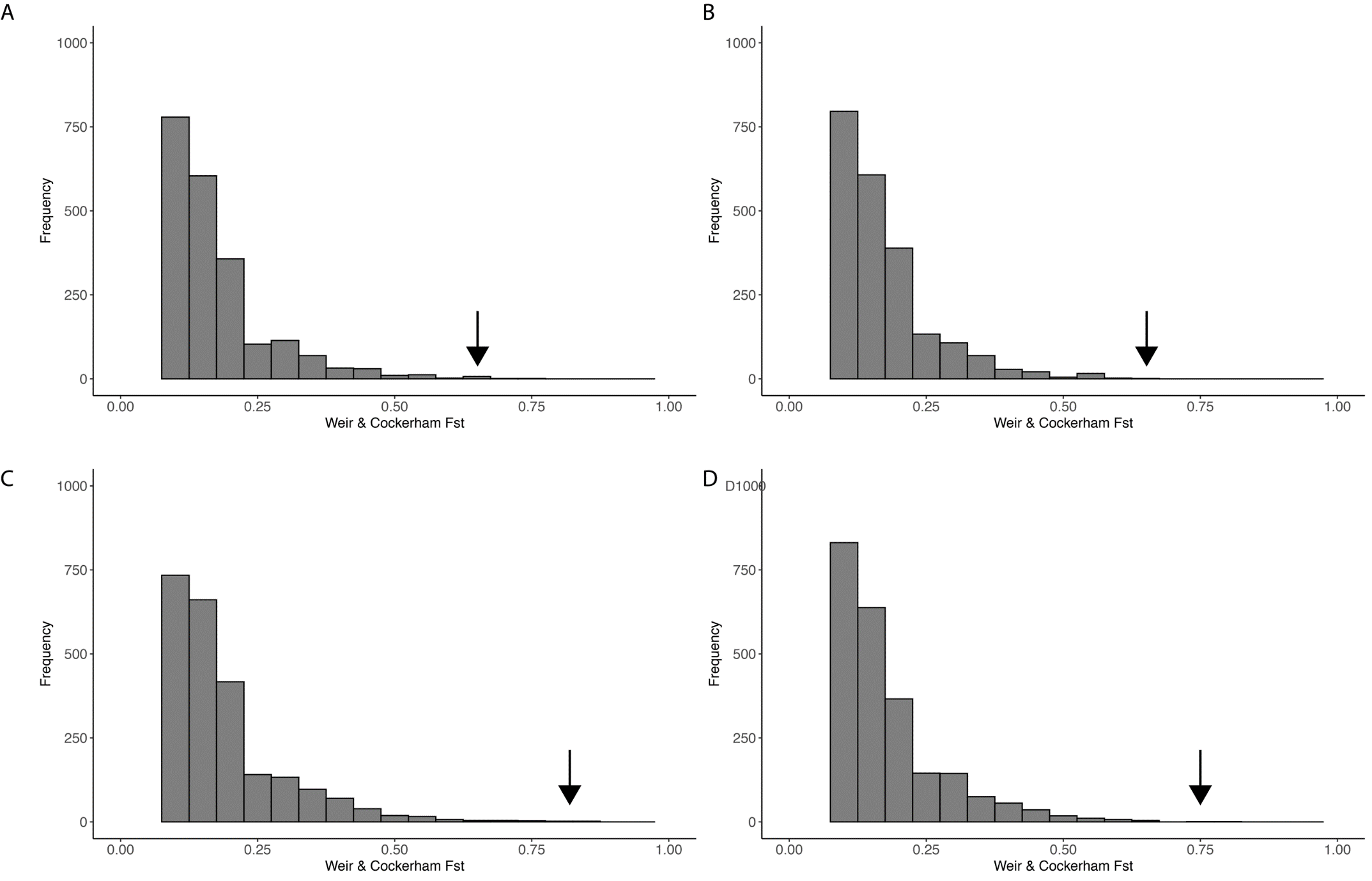
